# Tomato Protein Phosphatase 2C (SlPP2C3) negatively regulates fruit ripening onset and fruit gloss

**DOI:** 10.1101/2020.05.25.114587

**Authors:** Bin Liang, Yufei Sun, Juan Wang, Yu Zheng, Wenbo Zhang, Yandan Xu, Qian Li, Ping Leng

**Author notes:** These authors contributed equally to the present work (Bin Liang, Yufei Sun). Highlight: SlPP2C3 regulate tomato fruit ripening. To whom reprint requests should be addressed.

## Abstract

Abscisic acid (ABA) plays a vital role in coordinating physiological processes during fresh fruit ripening. ABA can bind to ABA receptors which interacts and inhibits their co-receptors type 2C phosphatases (PP2Cs). However, the dissected mechanism of PP2C during fruit ripening is unclear. In this study, we identify the role of SlPP2C3, a tomato type 2C phosphatase, as a negative regulator of ABA signaling and fruit ripening. SlPP2C3 selectively interacted with monomeric ABA receptors and SlSnRK2.8 kinase in both yeast and tobacco epidermal cells. Expressions of *SlPP2C3* were observed in all tissues, and it negatively correlated with the fruit ripening which was induced by exogenous ABA. Tomato plants with suppressed *SlPP2C3* expression exhibited enhanced sensitivity to ABA, while *SlPP2C3* over-expressed plants were less sensitive to ABA. Meaningfully, lack of *SlPP2C3* expression causes the acceleration of fruit ripening onset via the alternation of ABA signaling activity, and the fruit gloss is affected by the changes of outer epidermis structure. RNA-seq analysis found significant different expression of cuticle-related genes in pericarp between wild-type and *SlPP2C3* suppressed lines. Taken together, our finding demonstrate that SlPP2C3 plays an important role in the regulation of fruit ripening and fruit appearance quality in tomato.

## Introduction

Phytohormone ABA plays roles in various physiological processes during plant lifecycle, including seed dormancy and germination, root growth, stomata movement, leaf senescence and fruit ripening (Schroeder *et al.*, 2001; Finkelstein *et al.*, 2002; Zhang *et al.*, 2009a,b). Plant ABA response is largely relied on its signal transduction pathway (Cutler *et al.*, 2010). A dominant ABA signaling pathway are composed of PYR/PYL/RCAR family receptor proteins, group A type 2C protein phosphatases (PP2Cs), and subfamily 2 of the sucrose nonfermenting 1-related kinases (SnRK2s) (Fujii *et al.*, 2009; Ma *et al.*, 2009; Park *et al.*, 2009; Umezawa *et al.*, 2009; Vlad *et al.*, 2009). The canonical ABA signaling model could be summarized as following: ABA binding to PYR/PYL/RCAR receptors induces a conformational change of receptor and a compete SnRK2 kinase from PP2C inhibition. Released SnRK2s can be activated by Raf-like kinases, then downstream substrates can be phosphorylated, such as transcription factor or ion channel protein to elicit ABA response (Cutler *et al.*, 2010; Weiner *et al.*, 2010; Hauser *et al.*, 2011).

Protein phosphatases 2C is the earliest identified key component in ABA signaling pathway (Koornneef, 1984). *Arabidopsis* group A PP2C members ABI1 and ABI2 were cloned from *abi1-1* and *abi1-2* mutants with ABA-insensitive phenotypes, and were shown to play a negative role in ABA signaling regulation with overlapped functions (Leung *et al.*, 1994; Meyer *et al.*, 1994; Leung *et al.*, 1997; Gosti *et al.*, 1999; Merlot *et al.*, 2001). So far, nine genes encoded PP2C were identified as ABA signaling pathway components in *Arabidopsis*, including ABI1, ABI2, HAB1, HAB2, AHG1, AHG3/AtPP2CA, and three HAIs (Saez *et al.*, 2004; Kuhn *et al.*, 2006; Yoshida *et al.*, 2006; Nishimura *et al.*, 2007; Bhaskara *et al.*, 2012). PP2Cs interact and phosphorylate SnRK2s, the activity of PP2Cs is inhibited by ABA receptors PYR/PYL/RCAR proteins in both ABA-dependent and ABA-independent manners (Hao *et al.*, 2011; He *et al.*, 2014; Nemoto *et al.*, 2018). Structural studies indicate that Ser residues on the side chain of PYR/PYL/RCAR receptor insert into the PP2C active site and competitively occlude the access of the substrates (Melcher *et al.*, 2009; Miyazono *et al.*, 2009; Yin *et al.*, 2009). In this model, two residues are important for the interaction. One is Gly-180 of ABI1 or Gly246 of HAB1, this Gly residue is next to the PP2C active site and G160D mutation in ABI1 or Gly-Asp mutation in the corresponding position (Sheen, 1998; Robert *et al.*, 2006; Umezawa *et al.*, 2009; Yin *et al.*, 2009). Another important residue is Trp-300 of ABI1 or other corresponding Trp of the related PP2Cs which is the only residue in PP2C approaching ABA molecule in the interaction complex which plays an important role in complex stabilization (Melcher *et al.*, 2009; Dupeux *et al.*, 2011). Several studies show selective interaction among PP2C and PYR/PYL/RCAR receptors (Antoni *et al.*, 2012; Bhaskara *et al.*, 2012; Zhao *et al.*, 2013; Fuchs *et al.*, 2014), but the underlying mechanism remains unclear. Most of the PP2Cs have redundant function and show additive effect on ABA signaling. Single PP2C loss-of-function mutants exhibit mild phenotypic effects, while double or triple mutants show constitutive ABA response (Rubio *et al.*, 2009).

Although physiological function of PP2C function have been described in model plant *Arabidopsis* and rice (Saez *et al.*, 2004; Kuhn *et al.*, 2006; Yoshida *et al.*, 2006; Nishimura *et al.*, 2007; Bhaskara *et al.*, 2012), the function of PP2C on fresh fruit development and ripening is unknown. Recent studies suggested ABA played a role in metabolism in tomato fruit ripening regulation (Galpaz *et al.*, 2008; Zhang *et al.*, 2009; Sun *et al.*, 2012a; Sun *et al.*, 2012b; Sun *et al.*, 2017). However, seldom is known on the function of ABA signaling components during fresh fruit ripening. Overexpression of tomato ABA receptor gene *SlPYL9* leads to early fruit ripening, while *SlPYL9*-RNAi affects the mesocarp thickness and petal abscission (Kai *et al.*, 2019). In addition, the function of *SlPP2C1* was reported in tomato. *SlPP2C1* suppressed the early fruit ripening of plants but led to abnormal flower development, which may reduce the fruit yield (Zhang *et al.*, 2018).

To further understand ABA signaling during tomato fruit ripening, we identified the roles of SlPP2C3 in tomato fruit. It is found that SlPP2C3 may play a crucial role in fruit maturity and quality by affecting gene expressions of ethylene and cuticle.

## Materials and methods

### Plant materials

Tomato plants *(Solanum lycopersicum* L. cv. Micro Tom), including wild-type plants and transgenic lines, were grown under standard greenhouse conditions. For gene expression analysis, fruits were sampled according to the developmental stages, including mature green (MG), three days after breaker (B+3), ten days after breaker (B+10).

### Generation of SlPP2C3 transgenic tomato lines

For *SlPP2C3-OE* tomato transformation, *SlPP2C3* (Solyc06g076400.2.1) CDS was cloned into pRI101-AN plasmid (TAKARA). For *SlPP2C3*-RNAi tomato transformation, a fragment of the N-terminus specific sequence was amplified from the *SlPP2C3* cDNA and was cloned in both sense and anti-sense orientation into pCambia1305.1 plasmid to form a hairpin structure for RNA interference. The primers were shown in Table S2, in which *SlPP2C3-OE* and *SlPP2C3*-RNAi constructs were introduced into tomato (Micro Tom) via *Agrobacterium tumefaciens* LBA4404-mediated transformation. OE lines were identified through Kanamycin resistance and PCR detection. Positive *SlPP2C3*-RNAi transgenic plants were identified through hygromycin B resistance and GUS staining. Lines RNAi4 and RNAi13 were selected for further studies due to the suppression of *SlPP2C3* expression. T0 plants were self-pollinated to generate T1 and T2 plants. Phenotypic and molecular analyses of plants and fruits were performed on T2 plants after selfing of a single homozygote T1 plant, while drought resistance, seed germination and primary root growth assay were performed on T3 seeds obtained from selfing of homozygote T2 plants.

### Subcellular localization and bimolecular fluorescence complementation (BiFC) assay

To investigate the subcellular localizations of SlPP2Cs, the coding ORF sequences of *SlPP2Cs* tagged with GFP at the C-terminus were cloned into the pCambia1300 vector. The specific primers are listed in Table S3. *Agrobacterium tumefaciens* GV3101 carrying pCambia1300-SlPP2Cs-GFP was injected into the 5-6-week-old *Nicotiana benthamiana* leaves. 2 days after cultured in dark, the infiltrated leaves were examined, and the GFP signal was monitored by a fluorescence microscope (Olympus BX51, Japan) with an optical wavelength of 470 nm at 200x magnification.

For bimolecular fluorescence complementation, the cDNA of *SlPP2C3* was pSPYCE(M), and *SlPYLs* and *SlSnRK2s* were cloned into pSPYNE173 (Waadt *et al.*, 2008). The primers are listed in Table S3. *A. tumefaciens* GV3101 carrying pSPYCE(M) or pSPYNE173 fusion constructs were co-infiltrated into 5-6-week-old *N. benthamiana* leaves. After 2 days, the infiltrated leaves were examined via fluorescence microscope (Olympus BX51, Japan).

### Yeast two-hybrid (Y2H) assay

The full-length, site-directed mutagenic, and catalytic core (residues 105–411) of *SlPP2C3* were cloned into Y2H ‘prey’ vector pGADT7 (Clontech). Full-length SlPYLs and SlSnRK2s were cloned into the ‘bait’ vector pGBKT7 (Clontech). Prey and bait vectors were co-transformed into yeast strain Y2HGold (Clontech) and were grown on SD -Leu/-Trp plates at 30 °C for 3 days. The interactions were examined on the control media SD –Leu/-Trp and the selective media SD -Leu/-Trp/-His/-Ade in the presence or absence of 10 μM ABA. Yeast containing interest protein in combination with empty pGADT7 or pGBKT7 were used as controls. The primers used for Y2H assay are listed in Table S4.

### Site-directed mutagenesis

Mutants were created using the Fast Mutagenesis System (Transgen) according to the manufacturer’s instructions. Mutagenesis reactions were conducted with pGADT7-SlPP2C3 as template. All mutagenesis primer sequences are provided in Table S5.

### Seed germination and primary root growth assays

Wild-type and transgenic T3 generation seeds obtained from T2 homozygous plants, were harvested, air dried, and stored at 4□ for three weeks. After surface sterilization, the seeds (about 50 seeds for each replicate) were sown onto 1/2 MS medium containing 0 or 3 μM ABA. Then the seeds were germinated in the dark at 25°C for 4 or 10 d. Seed germination rates were investigated daily. Radicle emergence >1 mm indicated the success germination. Three replicate plates were used in each treatment. For primary root growth assay, wild-type, *SlPP2C3-OE,* and *SlPP2C3-RNAi* seedlings were grown on a 1/2 MS medium for 3 d, and then the seedlings with roots of the same length were transferred onto another 1/2 MS medium supplemented with 0 or 10 μM ABA and growth for 5 d. The root length was recorded from the base of the hypocotyl to the root tip. Experiments were performed three times.

### Drought stress treatment, measure of relative water content and detached leaf water loss

16-day-old WT and two independent lines of OE4 and RNAi13 were under a 22-day drought stress treatment by halting irrigation, and then the plants were subsequently re-watered for 5 days. The survival rate of plants was measured. Photographs were taken 0, 11, 15, 19, 22 d after the drought treatment and 5 d after re-water. The relative water contents (RWC) in the leaves of the plants were measured 0, 11, 15, 19, 22 days after the treatment. RWC was calculated by [(fresh weight – dry weight) / (turgid weight – dry weight)]. For the assay of water in detached leaves, healthy leaves were detached from each line and placed abaxial side up in an open plate under 25°C, 35% relative humidity condition. The water loss was measured over a period of 8 h. All experiments were repeated three times.

### qRT-PCR analysis

Total RNA was extracted from tomato tissues by the improved hot borate method (Wan and Wilkins, 1994). DNase treated RNA was used for cDNA synthesis through PrimeScript II 1st Strand cDNA Synthesis Kit (Takara) according to the manufacturer’s instructions. qRT-PCR was performed via SYBR Premix Ex Taq (Takara) and Rotor-Gene 3000 system (Corbett Research). qRT-PCR was conducted in three biological replicates, and each sample was analyzed at least thrice and was normalized using the *SAND* gene (Solyc03g115810) as an internal control. The relative fold expression for each gene was calculated by Rotor-Gene Q software using the delta Ct method. Primers used for the qRT-PCR analysis are provided in Table S6.

### Measurement of ABA and ethylene

The extraction and measurement of ABA was performed according to the description previously (Sun *et al.*, 2017). Briefly, three grams of tissues were ground and extracted by 80% (v/v) methanol at −20 □ for 18 h. After centrifugation, pellet was extracted by 80% methanol twice. The supernatant was dried under vacuum, and the residue was dissolved in a 0.02 M phosphate buffer (pH 8.0). Petroleum ether was used to remove pigments and the insoluble PVPP was used to remove polyphenols. ABA was extracted three times by ethyl acetate. The ethyl acetate phase was dried under vacuum and dissolved in 50% methanol (v/v). The ABA content was determined via HPLC (Agilent LC 1200, CA, USA) through a 4.8 x 150 mm C18 column (Agilent Technologies, CA, USA), and (±)-abscisic acid (Sigma) was used as standards. Three replicates were conducted for each sample.

Ethylene production was measured by enclosing two to four fruits in an 50-mL airtight containers for 2 h at 25°C, then 1 ml of the headspace gas was withdrew and injected into a gas chromatograph (Agilent GC 6890N, CA, USA) fitted with a flame ionization detector and an activated alumina column (Zhang *et al.*, 2009).

### RNA-seq

Total RNA was extracted from the fruit skin of WT and *SlPP2C3*-RNAi4 at mature green stage. RNA degradation and contamination were monitored on 1% agarose gels. RNA integrity was assessed through the RNA 6000 Pico Assay Kit of the Bioanalyzer 2100 system (Agilent Technologies, CA, USA). A total amount of 3μg RNA per sample was used for mRNA purification and library construction via the Truseq™ RNA Sample Prep Kit (Illumina, CA, USA) followed the manufacturer’s instructions. The samples were sequenced on an Illumina HiSeq™ 2000 (Illumina). Each sample yielded more than 6 G of data.

Raw data of fastq format was firstly processed through in-house perl scripts. Clean data was obtained by removing reads containing adapter, reads containing ploy-N and low-quality reads from raw data. Filtered RNA-seq clean reads were aligned to the reference tomato genome releasing SL2.50 through TopHat v2.0.13 (Kim *et al.*, 2013). HTSeq v0.5.3 was used to count the reads numbers mapped to each gene. And then the RPKM of each gene was calculated based on the length of the gene and reads count mapped to this gene. Differential expression analysis was performed using the DEGSeq R package (1.12.0) and Cufflinks (Trapnell *et al.*, 2012). Corrected P-value of 0.001 and log2 (Fold change) of 1 were set as the threshold for significant differential expression.

### Statistical analysis

The data were statistically analyzed with SPSS software using one-way analysis of variance and Duncan’s test of significance. *t-test P value < 0.05; **t-test P value < 0.01.

## Results

### Phylogenetic analyses of tomato Group A PP2C family

To explore the entire family members of group A PP2C in tomato, predicted *Arabidopsis* PP2C protein sequence (Schweighofer *et al.*, 2004) was used as query to BLAST against the SGN database (https://solgenomics.net/). Total 93 putative protein phosphatase 2C family members were identified in tomato genome. Phylogenetic analyses indicated that tomato PP2Cs were divided into 12 groups (A-K) (Fig. S1), which was consistent with previous study in *Arabidopsis* and rice (Schweighofer *et al.*, 2004; Xue *et al.*, 2008). Fourteen tomato PP2Cs were identified as members of group A, which have been shown to participate in ABA signaling. We named these 14 tomato members as SlPP2C1-14 (Table S1). Further amino acid alignment indicated the conserved catalyze domain of 14 PP2Cs with *Arabidopsis* homologues and diverse N-terminal (Fig. S2). All tomato PP2Cs have conserved active-site residues and PYL-interaction residues, except SlPP2C7 in which some motifs in catalyzed domain is missing. Interestingly, SlPP2C11-14 lack the conserved tryptophan residue which acts as a “lock” in the interaction with ABA receptor, similar to *Arabidopsis* AtAHG1 (Antoni *et al.*, 2012). It is worth noting that SlPP2C3 is the single ortholog of *Arabidopsis* AtHAI1/2/3 (Fig. S3). In *Arabidopsis,* three AtHAIs have redundant function during drought resistance, leaf senescence and seed dormancy (Bhaskara *et al.*, 2012; Zhang *et al.*, 2012; Kim *et al.*, 2013) which implied the non-redundant function of SlPP2C3 in tomato physiological processes.

### SlPP2C3 selectively interact with monomer ABA receptors and SlSnRK2.8

Our previous study reported the interactions of SlPP2C1-5 and SlPYLs or SlSnRK2s which showed that SlPP2C3 is the only member interacted with SlSnRK2 in yeast two hybrid assay (Chen *et al.*, 2016). In this work, we further study the interaction manner of SlPP2C3 and ABA receptors. Full length of SlPP2C3 interacted with subfamily I and subfamily II monomeric receptors in yeast cell. The interactions between SlPP2C3 and SlPYL1, 4, 7, 9 or 11 was ABA dependent while the interactions with SlPYL2, 3, 10 or 13 were observed as ABA independent. In addition, SlPP2C3 did not interact with subfamily III dimeric receptor members SlPYL5 and 8 even in the presence of 10 μM ABA (Fig. 1A). For SlSnRK2s, SlPP2C3 can only interact with SlSnRK2.8 (Fig. 1B), consistent with our previous results (Chen *et al.*, 2016).

**Fig. 1.**
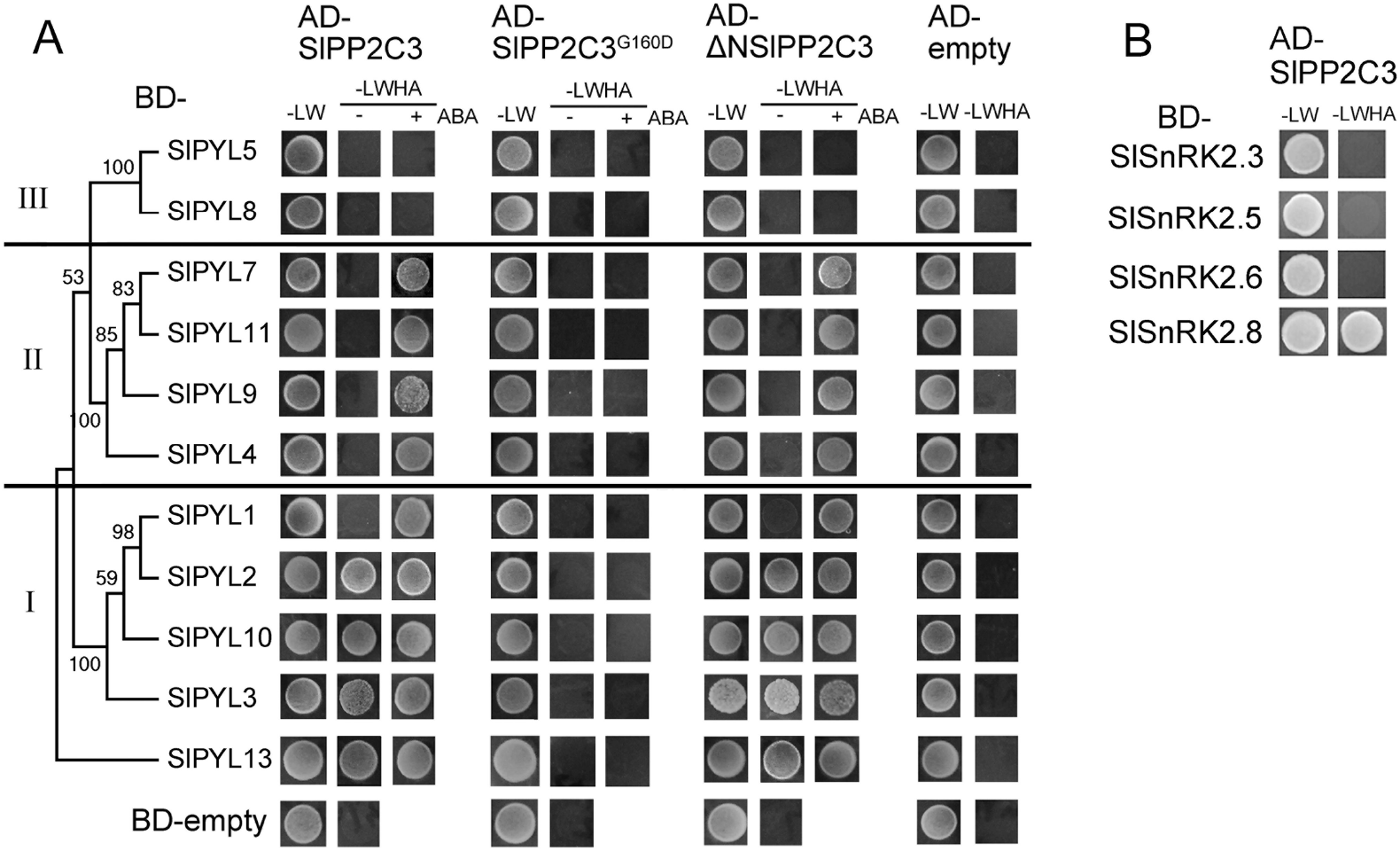
Interactions of SlPP2C3 with SlPYLs or SlSnRK2s in yeast two hybrid (Y2H) assay. Interactions were determined using a growth assay on media lacking Leu, Trp, His and Ade (-LWHA), with or without 10 μM ABA. The yeasts growth on media lacking Leu and Trp (-LW) were used as control. (A) Interaction of SlPP2C3-AD and SlPYLs-BD receptors. First lane, full length SlPP2C3; second lane, SlPP2C3 with Gly-160 mutated into Asp (SlPP2C3^G160D^); third lane, the catalytic core (residues 105-411) of SlPP2C3 (ΔNSlPP2C3); last lane, pGADT7 empty vector control. (B) Interaction of SlPP2C3-AD and SlSnRK2s-BD.

Catalyzed domain of SlPP2C3 (105-411 amino acid) acted similar as the full length. However, the Gly-160Asp (corresponding to AtABI1^G180D^ encoded by *abi1-1)* mutation abolished the interaction between SlPP2C3 and all SlPYLs (Fig. 1A). Further, bimolecular fluorescence complementation (BiFC) assay was performed to confirm the SlPP2C3-SlPYLs/SlSnRK2.8 interaction *in-planta.* Fluorescent signal was detected in tobacco leaf which co-expressed in SlPP2C3 and SlPYL1, 3, 9, 10, 13 or SlSnRK2.8 (Fig. 2A). The interaction between SlPP2C3 and SlPYL1/9 may due to the sufficient endogenous ABA level in leaf. Most interactions occur in nucleus and cytoplasm, except for SlPP2C3 and SlPYL3, which only occur in nucleus. The interaction localization was further validated by subcellular location of SlPP2C3 (Fig. 2B). Both *in-vitro* and *in-planta* assay indicate that SlPP2C3 cannot interact with dimer receptors belonging to subfamily III (Fig. 1A; Fig. 2A).

**Fig. 2.**
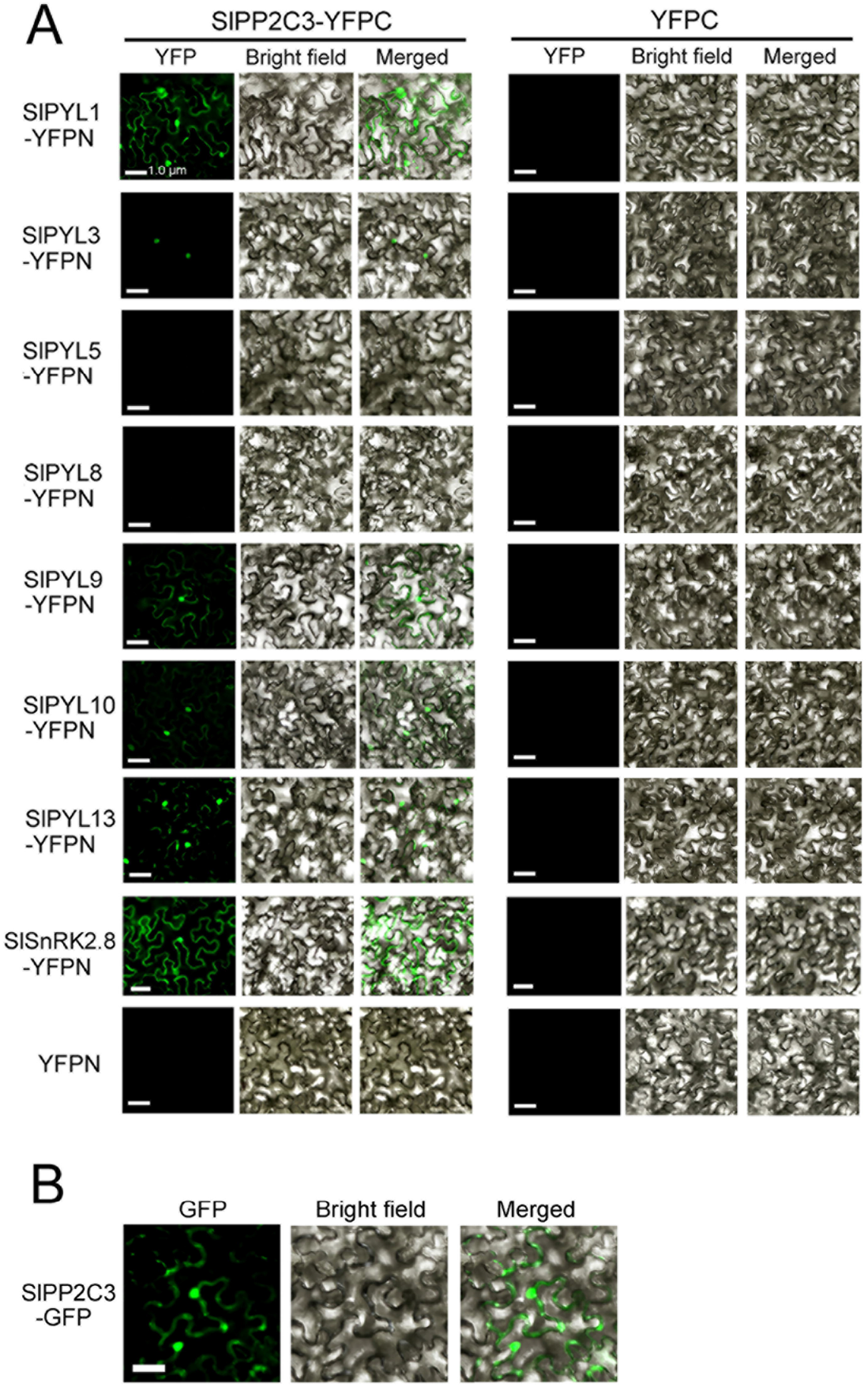
Interaction of SlPP2C3 with SlPYLs and SlSnRK2s in tobacco leave. (A) SlPP2C3-YFPC and SlPYLs-YFPN or SlSnRK2s-YFPN were co-expressed in *N. benthamiana* leaves through *A. tumefaciens* infiltration. The YFP signal was monitored 2 days after co-infiltration. (B) Subcellular locations of SlPP2C3. SlPP2Cs-GFP was transiently expressed in *N. benthamiana* leaves through *A. tumefaciens* infiltration. The GFP signal was monitored 2 days after infiltration. The left column shows fluorescent signal; the middle column shows the bright-field image; and the right column shows the merged image.

Our previous study showed that subfamily III receptors SlPYL5 and SlPYL8 interacted with SlPP2C5, ortholog of *Arabidopsis* HAB1. To understand molecular mechanism of these selective receptor interaction, we compared the amino acid sequences of SlPP2C3 and SlPP2C5, and found some diverse amino acid residues in PP2C-PYL interaction region between SlPP2C3 and SlPP2C5 (Fig. 3A) (Melcher *et al.*, 2009; Miyazono *et al.*, 2009). We conducted the site-mutagenesis on SlPP2C3 to switch these residues to the SlPP2C5 version, reasoning that the mutant of these residues could mimic SlPP2C5 (Fig. 3A). Our results revealed that S295F mutation in SlPP2C3 increased the interactions of SlPP2C3-SlPYL5/8, while R285K mutation increased SlPP2C3-SlPYL8 interaction in the presence of ABA (Fig 3B). These results partially explained the preferred interaction of SlPP2Cs and SlPYLs.

**Fig. 3.**
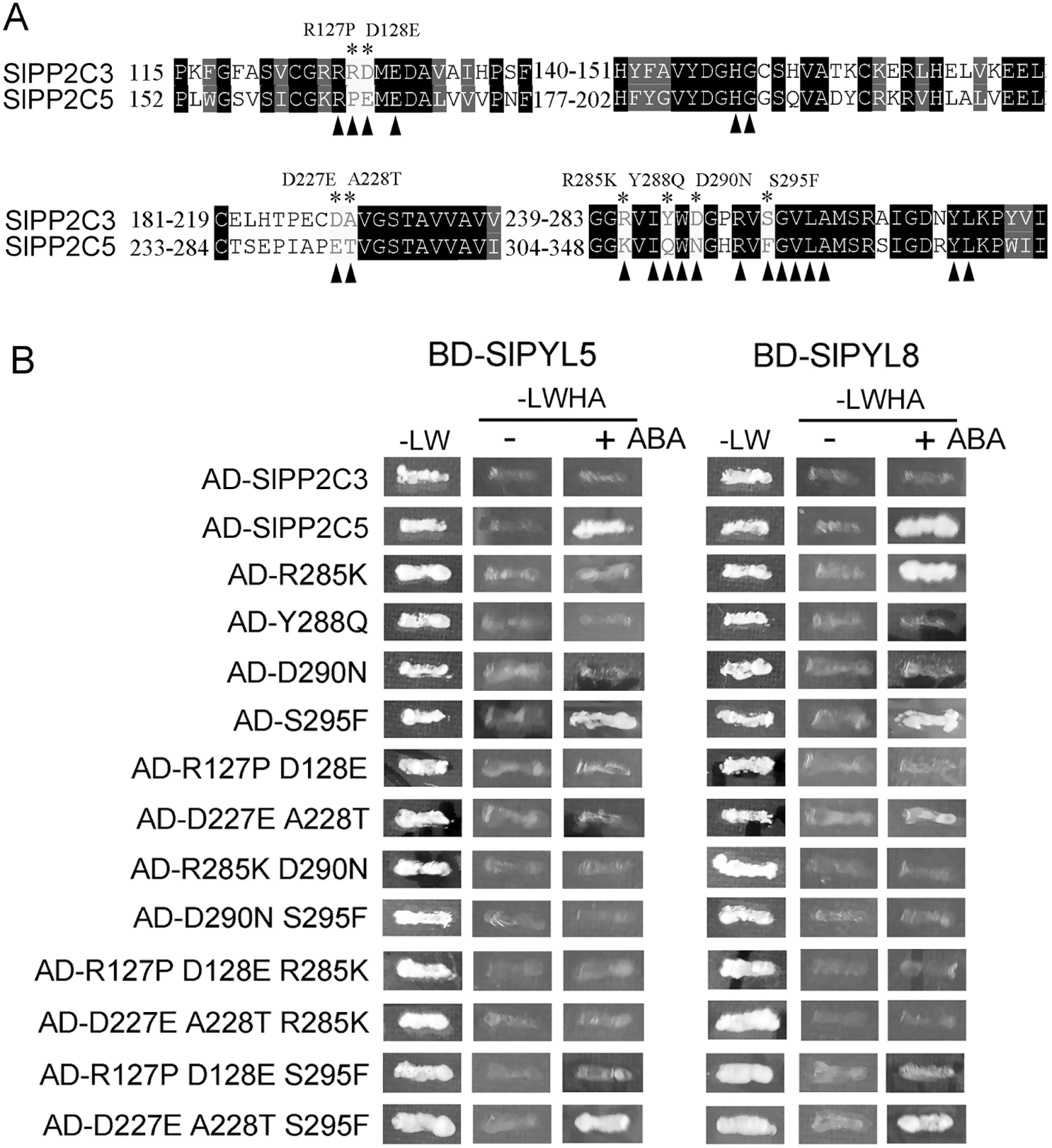
*SlPP2C3* mutations interact with dimeric receptors SlPYL5 and SlPYL8 in Y2H assay. (A) Alignment of SlPP2C3 and SlPP2C5 amino acid sequences. Location of the SlPP2C3 mutations are marked by black asterisks. Black triangle indicated the residues involved in PYLs interaction (Miyazono *et al.*, 2009). (B) Interaction of SlPP2C3, SlPP2C5 or SlPP2C3 mutants (fused to the activating domain) and either SlPYL5 or SlPYL8 (fused to the binding domain). Interaction was determined by growth assay on medium lacking Leu, Trp, His and Ade (-LWHA), with or without 10 μM ABA. The photographs were taken after 5 d growth at 30.□

### SlPP2C3 expression is negatively correlated with fruit ripening and induced by ABA

*SlPP2C3* was highly transcribed in seed, however, the expression in seedling, root and stem were relatively low. During flowering and fruit development, *SlPP2C3* mRNA level was high in flower and immature fruit and it decreased during ripening, which indicated that *SlPP2C3* might play a negative role in fruit ripening (Fig. 4A).

**Fig. 4.**
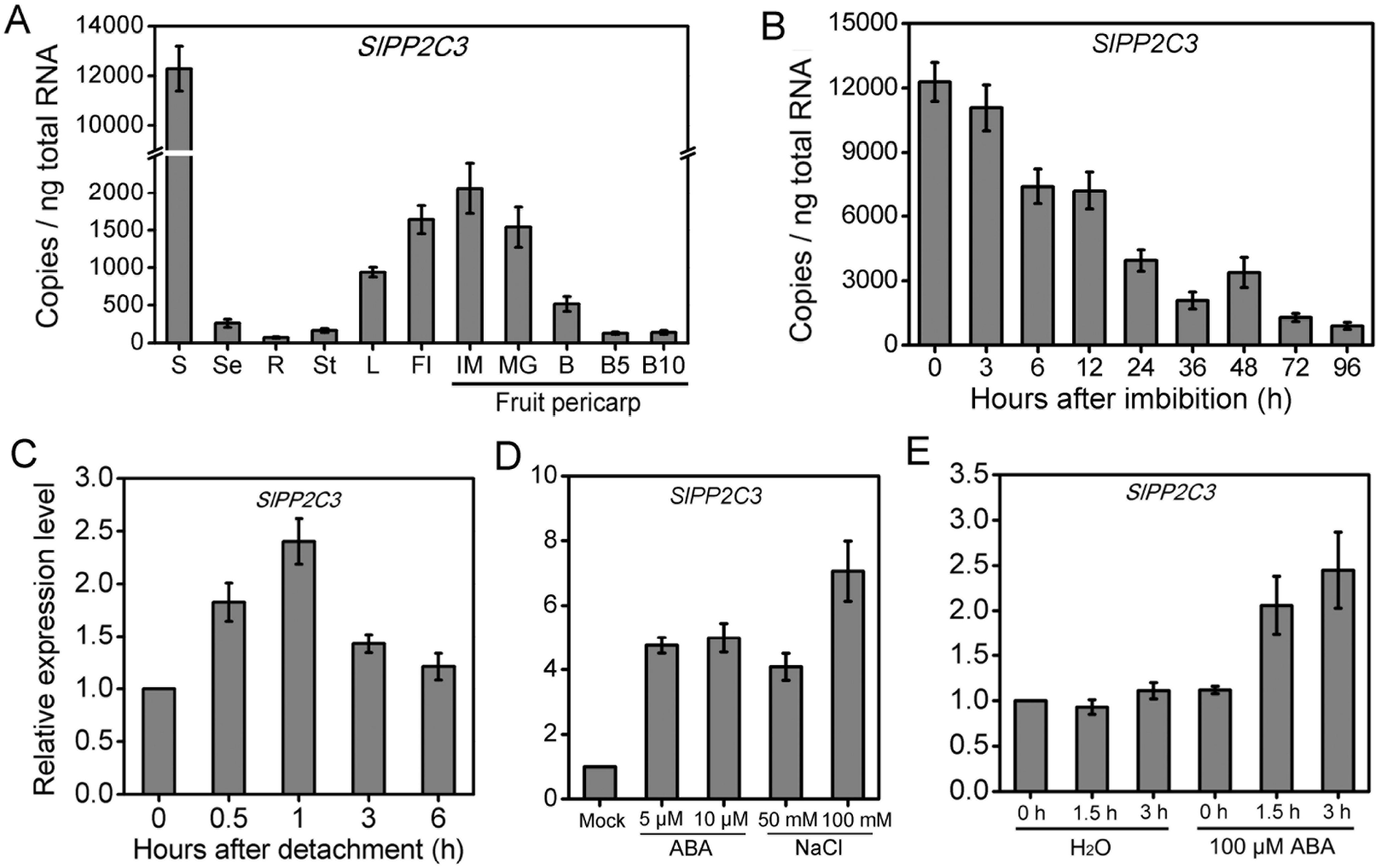
Expression pattern of *SlPP2Cs* genes in tomato. (A) Absolute quantification of *SlPP2C3* in various tomato tissues. S, seed; Se, seedling; R, root; St, stem; L, Leaf; Fl, flower; fruit pericarp at IM (immature, 15 DAF), MG (mature Green, 25 DAF), B (break), B5 (5 days after break) and B10 (10 days after break). Error bars represent ± SD of biological triplicates. (B) Absolute quantification of *SlPP2C3* in tomato seeds which were soaked and imbibed in water for indicated time. (C) Relative expression of *SlPP2C3* in tomato detached leaves. (D) Relative expression of *SlPP2C3* in tomato seeds which were treated with exogenous ABA or NaCl for 48 h. (E) Relative expression of *SlPP2C3* in mature green tomato pericarp disks treated by 100 μM exogenous ABA. For all expression analysis, *SlSAND* gene was used as an internal control. The error bars indicate ± SD of biological triplicates. *P value t-test < 0.05; **P value t-test < 0.01.

To examine whether *SlPP2C3* transcription is regulated by ABA, we performed qRT-PCR to investigate the expressions of *SlPP2C3* during seed imbibition and leaf dehydration. These two physiological processes have been shown to accompany by the decreasing or increasing of endogenous ABA level, respectively. The expression of *SlPP2C3* decreased during seed imbibition, on the other hand, *SlPP2C3* was induced in dehydrated leaf (Fig. 4B-C). Furthermore, *SlPP2C3* were induced by exogenous ABA treatment in seed and mature green fruit (Fig. 4D-E). These results indicated that the expressions of *SlPP2C3* is positively regulated by ABA.

### Altered expression of SlPP2C3 resulted in the change of ABA sensitivity

Considering the expression and interaction pattern, we are interested in the physiological function of SlPP2C3 *in planta.* To gain insight into the role of *SlPP2C3,* the expressions of *SlPP2C3* through RNA interference were suppressed or overexpressed in Micro-Tom tomato, respectively. Independent *SlPP2C3-OE* (OE4 and OE33) and *SlPP2C3*-RNAi (RNAi4 and RNAi13) transgenic lines with dramatically increased or decreased *SlPP2C3* expression were used for further investigation (Fig. S4).

*SlPP2C3-OE* and *SlPP2C3*-RNAi plants were similar to WT in plant architecture (Fig. S5A-D). The leaves of transgenic lines exhibited similar shape, color and size to WT (Fig. S5F-G). These data suggested that SlPP2C3 played a limited role in plant vegetative growth.

SlPP2C3 is a putative core component in ABA signaling, therefore, we tested whether ABA sensitivity can be altered by *SlPP2C3* manipulation. The ABA-mediated inhibition of seed germination was compared among the transgenic lines and WT. It is found that the seed germination of *SlPP2C3*-RNAi lines was delayed which is more sensitive to exogenous ABA, while *SlPP2C3-OE* lines exhibited oppositely (Fig. 5A-B). Furthermore, the primary root growth of *SlPP2C3-OE* seedling was less sensitive to exogenous ABA treatment compared to WT. However, *SlPP2C3*-RNAi seedling was more sensitive to exogenous ABA treatment (Fig. 5C-D). These results demonstrated that *SlPP2C3* might act as a negative regulator in ABA signaling.

**Fig. 5.**
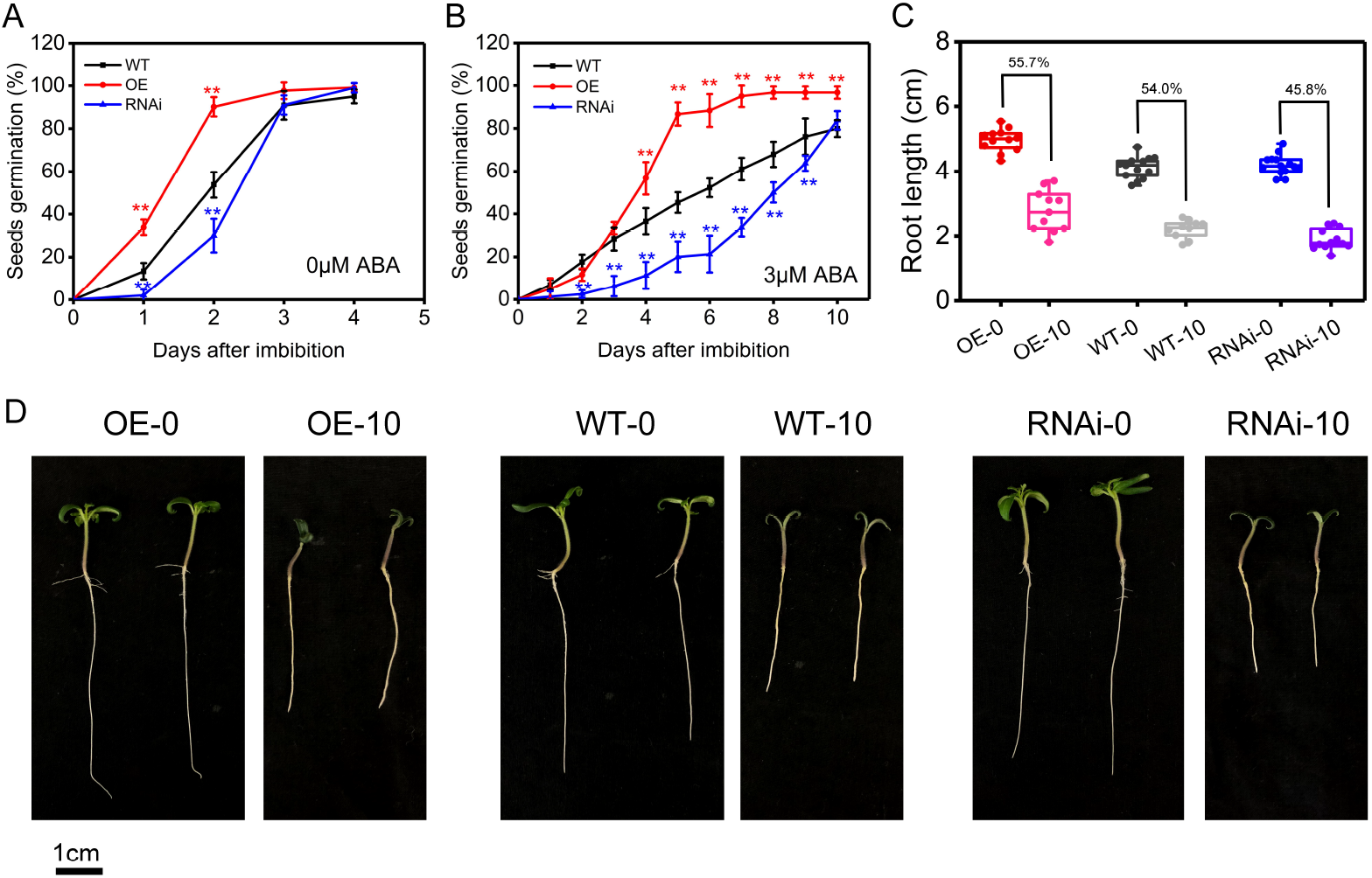
Altered sensitivity to ABA-mediated inhibition of seed germination, primary root growth in *SlPP2C3* transgenic lines compared to WT. (A, B) Approximately 50 seeds of *SlPP2C3-OE, SlPP2C3*-RNAi lines and WT (three independent experiments) were sown on 1/2 MS medium with or without 3 μM ABA. Seeds germination were scored every day. Values are means ± SE. *t-test P value < 0.05; **t-test P value < 0.01. (C, D) Primary root length and photographs of tomato seedlings 5 days after transferring 3-day-old seedlings into 1/2 MS medium with or without 10 μM ABA. OE: OE4 seeds mixed with OE33 seeds. RNAi: RNAi4 seeds mixed with RNAi13 seeds. OE-0/OE-10: OE seedlings 5 days after transferring 3-day-old seedlings into 1/2 MS medium with or without 10 μM ABA. Percentages represent the ratio of root length of seedlings treated with 10 μM ABA to that treated wuthout ABA. Values are means ± SD. *t-test P value < 0.05; **t-test P value < 0.01.

### SlPP2C3 plays a negative role in drought tolerance

To understand whether *SlPP2C3* plays a role in ABA-mediated drought tolerance, 16-days-old transgenic lines and WT plants were exposed to a 22-day drought stress treatment. 15 d after the treatment, *SlPP2C3-OE* line showed more wilted leaves than WT, while *SlPP2C3*-RNAi exhibited significant improvement under drought tolerance compared to the WT (Fig. 6A). 5 days after re-watering, *SlPP2C3-OE* plants showed 6% survival rate compared to 30% for the WT, while the survival rate of *SlPP2C3-*RNAi plants was nearly 60% (Fig. 6B). Consistent with the above result, the relative water content in *SlPP2C3-OE* leaves was significantly lower than that of WT, while the relative water content in *SlPP2C3-*RNAi leaves were the highest (Fig. 6C). Leaves from *SlPP2C3-*RNAi exhibited lower water loss rate than WT which implied the lower transpiration rate of *SlPP2C3-*RNAi leaves (Fig. 6D). These assays suggested that SlPP2C3 played a negative role in plant drought tolerance.

**Fig. 6.**
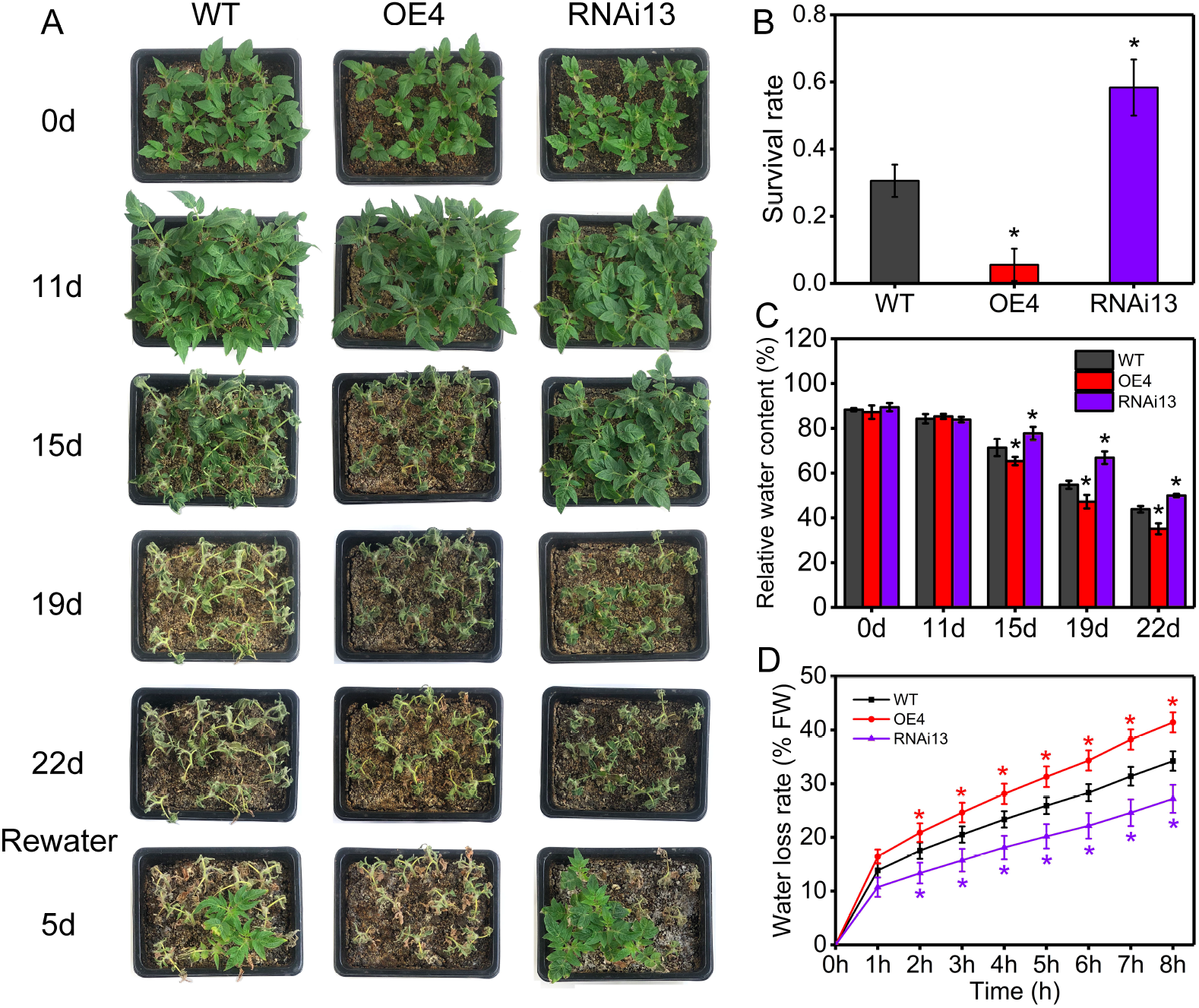
Drought resistance of *SlPP2C3* transgenic lines. (A) 16-days-old *SlPP2C3* transgenic lines and WT plants were subjected to drought stress withholding water for 22 days and then re-watered for 5 days. The experiments were repeated three times. (B) Survival rates were scored 5 days after re-watering. (C) Relative water contents in *SlPP2C3* transgenic lines and WT leaves after drought stress. Values are means ± SE. *t-test P value < 0.05; **t-test P value < 0.01. (D) Water loss analysis for the detached leaves. Values are means ± SD for three independent experiments. *t-test P value < 0.05; **t-test P value < 0.01.

### SlPP2C3 affects fruit ripening schedule

Given the role of ABA in fruit ripening regulation and the negative correlated expression pattern of *SlPP2C3* with fruit ripening, we investigated whether *SlPP2C3* played a role in fruit ripening. It was observed that the *SlPP2C3-OE* fruits required two more days from anthesis to the breaker stage than WT, while the *SlPP2C3*-RNAi fruits required 2-3 fewer days (Fig. 7A-B). The breaker time of WT is from 37 to 42 days, while the breaker times of *SlPP2C3-OE* and *SlPP2C3*-RNAi fruits are 39-44 days and 34-40 days, respectively. The ethylene and ABA production peaks of *SlPP2C3-OE* / *SlPP2C3*-RNAi fruits were delayed / advanced (Fig. 7C-D). Moreover, the transcription of some 1-aminocyclopropane-1-carboxylic acid oxidase genes which involved in ethylene biosynthesis and several ethylene receptor genes were up-regulated in *SlPP2C3*-RNAi fruits (Fig. S6; Fig. 10B).

**Fig. 7.**
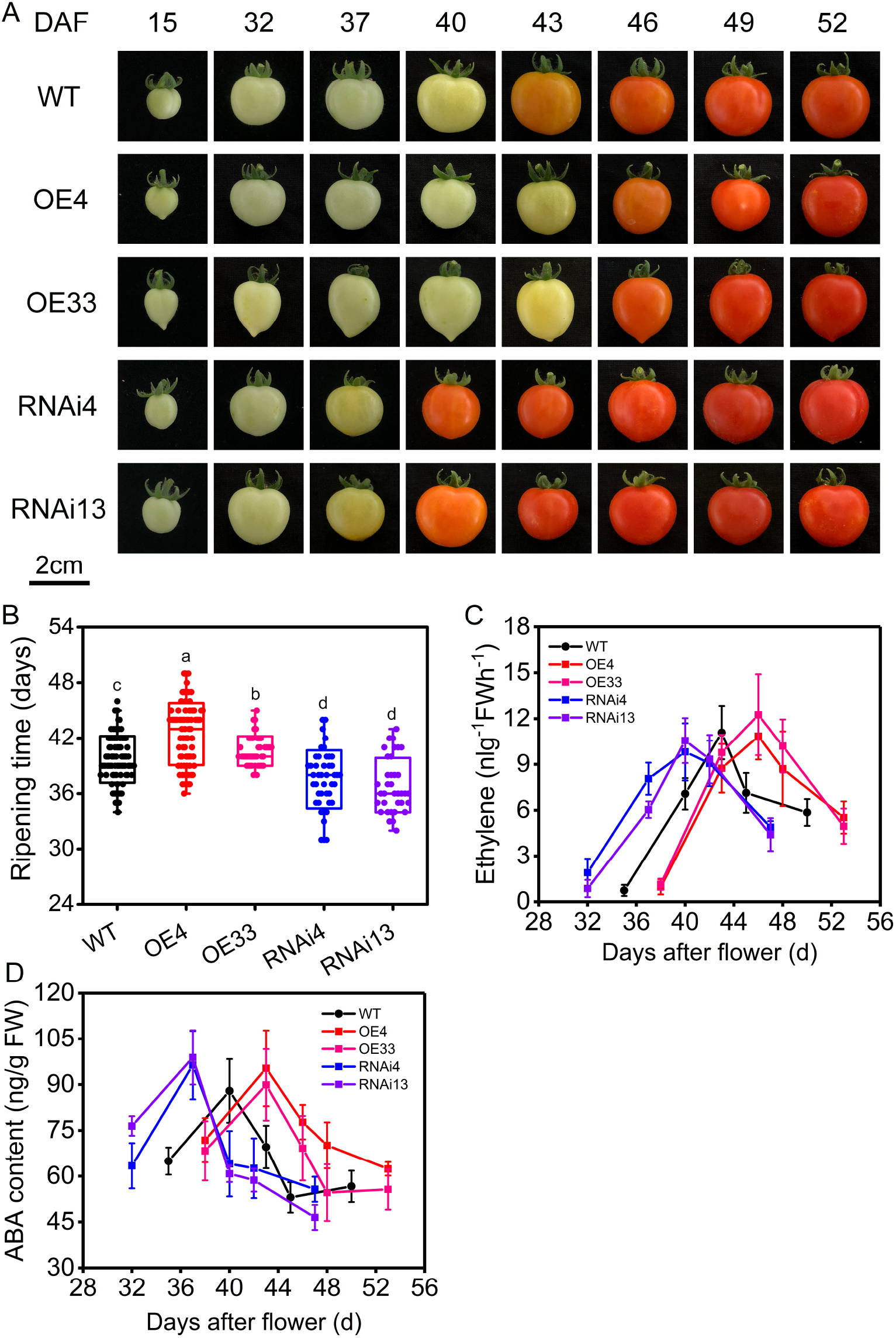
The phenotype of *SlPP2C3* transgenic and WT fruits. (A) The ripening time of fruits from *SlPP2C3* transgenic lines compared to WT fruits. (B) Days from anthesis to break. (C-E) Changes of ethylene release and ABA content in WT and *SlPP2C3* transgenic fruits during fruit ripening. n > 15, values are means ± SD. *t-test P value < 0.05; **t-test P value < 0.01.

To determine whether fruit quality was altered by *SlPP2C3-OE* or *SlPP2C3*-RNAi, several typical fruit quality parameters were measured. The fruit shape index (the ratio of fruit vertical diameter to horizontal diameter) of *SlPP2C3-OE* (especially OE33) was higher than WT (Fig. 8C). The fruit weight and seed numbers of *SlPP2C3-OE* were lower than WT, while *SlPP2C3*-RNAi showed no significant difference (Fig. 8D, F). Fruit firmness, contents of soluble solids, total sugar content and the titratable acid in the transgenic fruits were similar to WT (Fig. 8E, H; Fig. S7D-E). The number of OE33 fruits in 2 locule of OE33 was significantly higher than that of WT, while the number of OE33 fruits in 3 locule was less (Fig. 8G). The total carotenoids and lycopene of *SlPP2C3-OE* fruits were higher than WT, however, the *SlPP2C3*-RNAi fruit showed similar level to WT (Fig. S7A-C). These results suggest that SlPP2C3 play a role in fruit developed and metabolic processes.

**Fig. 8.**
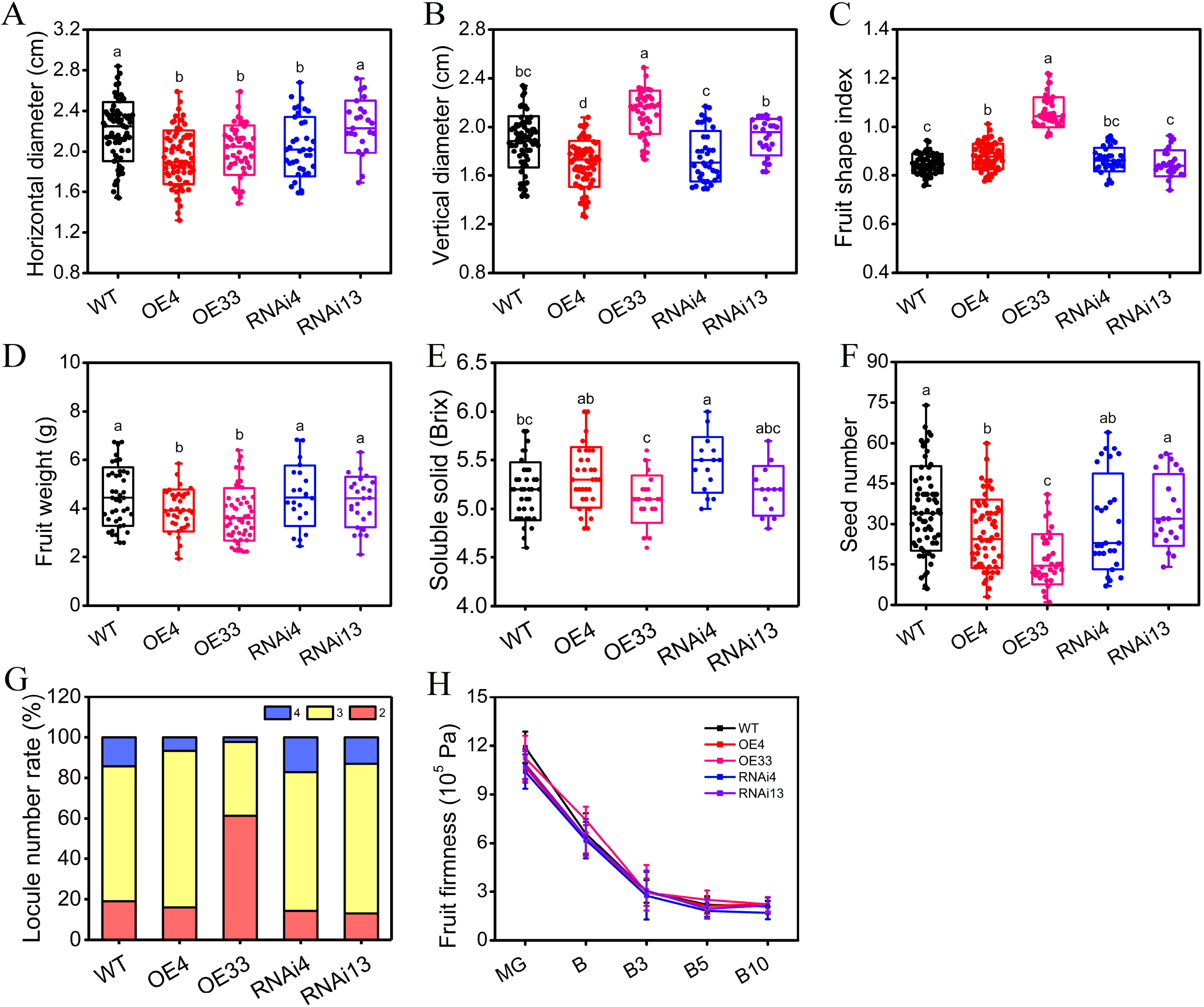
Physiological parameters related to fruit in *SlPP2C3* transgenic and WT fruits. (A) Horizontal diameters of B10 fruits. (B) Vertical diameters of B10 fruits. (C) Fruit shape index of B10 fruits. (D) Single fruit weight of B10 fruits. (E) Soluble solid content (Brix) of B10 fruits. (F) Seed number of each fruit. (G) Locule number rate of fruit. (H) Fruit firmness during fruit ripening. Values are means ± SD. Letters represent significant differences, t-test P value < 0.05.

### SlPP2C3-RNAi led to a bad fruit gloss due to altered outer epidermis

It is noticed that *SlPP2C3*-RNAi fruits exhibit dull fruit surface compared to the WT (Fig. 9A). Further scanning electron microscope study revealed a verrucous outer epidermis surface of *SlPP2C3*-RNAi fruit, which may imply an alter in cuticle/wax metabolism (Fig. 9B). To understand the underlying mechanism, a global gene expression analysis of *SlPP2C3*-RNAi fruit skin was performed (Dataset S1). 374 up-regulated and 519 down-regulated genes of *SlPP2C3*-RNAi fruit skin were screened and compared to the WT (Dataset S2). Among them, two transcription factors and 24 structural genes related to cuticle formation were identified (Table 1), and the expressions of some genes were further verified by qRT-PCR (Fig. 9D). R2R3-type MYB transcription factor *SlMYB96* (Solyc03g116100) were down-regulated in *SlPP2C3*-RNAi fruits, while *SlMYB106* (Solyc02g088190) were up-regulated. The orthologs in *Arabidopsis* were shown to regulate gene expressions in cutin and wax biosynthesis (Seo *et al.*, 2011; Oshima *et al.*, 2013). Out of 18 up-regulated genes, 11 were involved in cutin biosynthesis, including one *LACS* gene *SlLACS2* which was putatively required for the acyl activation of fatty acid (Schnurr *et al.*, 2004), four *CYP450s* genes *(SlCYP77A1, SlCYP77A2, SlCYP86A68* and *SlCYP86A69)* which were involved in midchain and terminal hydroxylation of fatty acid (Li-Beisson *et al.*, 2009; Shi *et al.*, 2013), four *SlHTH* genes in which the ortholog in *Arabidopsis* was proved to be important in the formation of dicarboxylic fatty acids (Kurdyukov *et al.*, 2006) and two Glycerol-3-phosphate acyltransferase genes *SlGAPT4* and *SlGAPT6* were mediated in the last step of cutin monomers biosynthesis (Li *et al.*, 2007), a cuticle transporter gene *SlABCG32,* and a cutin synthase gene *SlCD1* which was required for cutin polymerization (Bessire *et al.*, 2011; Yeats *et al.*, 2014; Fabre *et al.*, 2016). On the other hand, 7 out of 9 down-regulated genes were involved in wax biosynthesis. These data indicated that low *SlPP2C3* expression may change the cuticle composition of fruit outer epidermis.

**Fig. 9.**
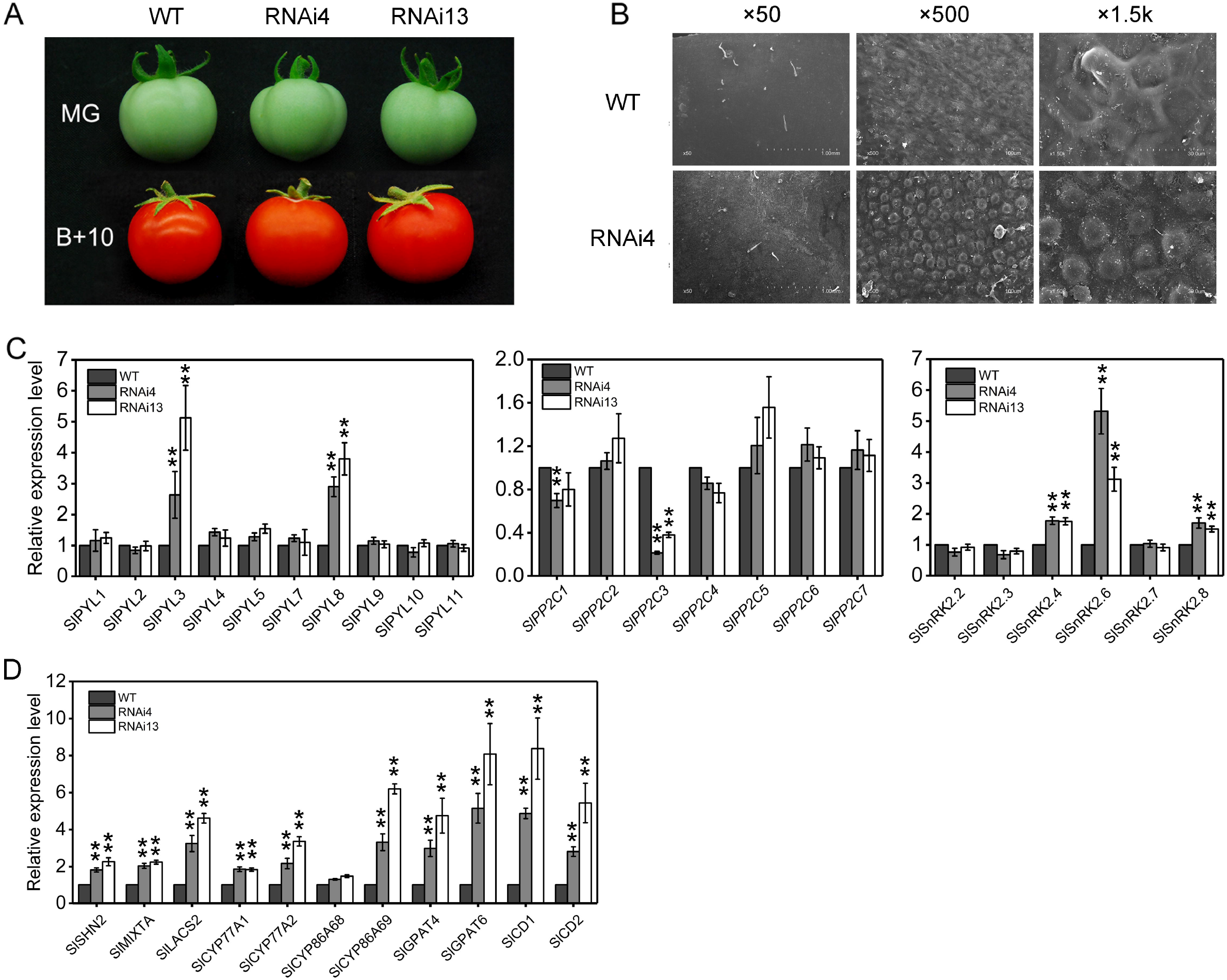
Reduced fruit brightness and altered gene expressions in cuticle metabolism and ABA signaling of *SlPP2C3*-RNAi fruits skin. (A) Photographs of *SlPP2C3*-RNAi lines and WT tomato fruit at MG and B+10 stages showing surface brightness. (B) Outer epidermis surface of *SlPP2C3*-RNAi lines and WT tomato fruits at B stage by scanning electron microscope. (C) Expressions of cuticle metabolism genes in transgenic and WT fruit skin at the MG stage. (D) Expressions of ABA signaling genes *(SlPYLs, SlPP2Cs* and *SlSnRK2s)* in transgenic and WT fruit skin at MG stage. *SlSAND* gene was used as internal reference. The data indicates the mean ± SD of three biological replicates. *t-test P value < 0.05; **t-test P value < 0.01.

**Table 1.**
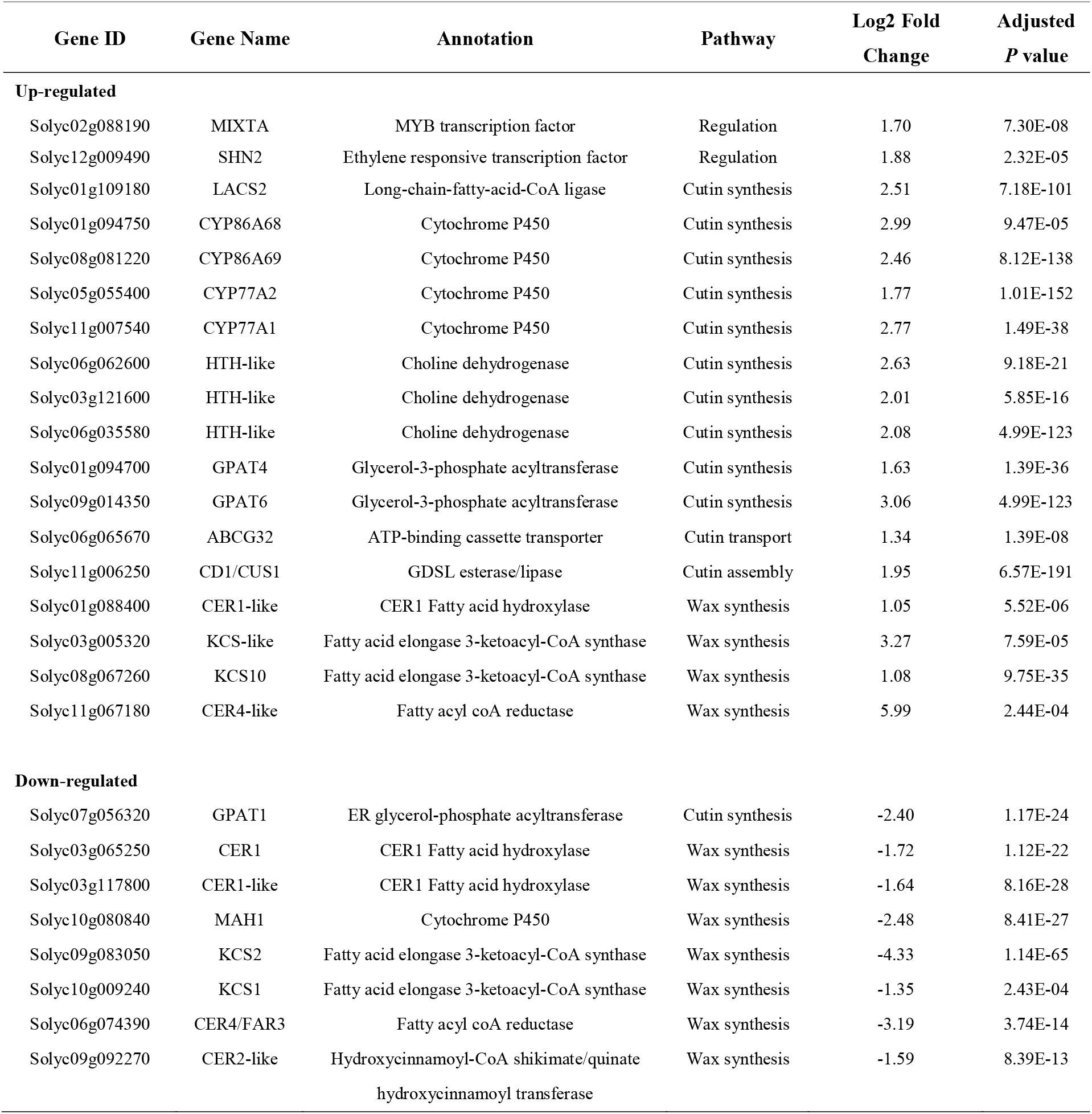
Differential expressed genes related to cuticle metabolism in transgenic fruit peel compared with WT.

| Gene ID | Gene Name | Annotation | Pathway | Log2 Fold Change | Adjusted *P* value |
| --- | --- | --- | --- | --- | --- |
| **Up-regulated** |  |  |  |  |  |
| Solyc02g088190 | MIXTA | MYB transcription factor | Regulation | 1.70 | 7.30E-08 |
| Solyc12g009490 | SHN2 | Ethylene responsive transcription factor | Regulation | 1.88 | 2.32E-05 |
| Solyc01g109180 | LACS2 | Long-chain-fatty-acid-CoA ligase | Cutin synthesis | 2.51 | 7.18E-101 |
| Solyc01g094750 | CYP86A68 | Cytochrome P450 | Cutin synthesis | 2.99 | 9.47E-05 |
| Solyc08g081220 | CYP86A69 | Cytochrome P450 | Cutin synthesis | 2.46 | 8.12E-138 |
| Solyc05g055400 | CYP77A2 | Cytochrome P450 | Cutin synthesis | 1.77 | 1.01E-152 |
| Solyc11g007540 | CYP77A1 | Cytochrome P450 | Cutin synthesis | 2.77 | 1.49E-38 |
| Solyc06g062600 | HTH-like | Choline dehydrogenase | Cutin synthesis | 2.63 | 9.18E-21 |
| Solyc03g121600 | HTH-like | Choline dehydrogenase | Cutin synthesis | 2.01 | 5.85E-16 |
| Solyc06g035580 | HTH-like | Choline dehydrogenase | Cutin synthesis | 2.08 | 4.99E-123 |
| Solyc01g094700 | GPAT4 | Glycerol-3-phosphate acyltransferase | Cutin synthesis | 1.63 | 1.39E-36 |
| Solyc09g014350 | GPAT6 | Glycerol-3-phosphate acyltransferase | Cutin synthesis | 3.06 | 4.99E-123 |
| Solyc06g065670 | ABCG32 | ATP-binding cassette transporter | Cutin transport | 1.34 | 1.39E-08 |
| Solyc11g006250 | CD1/CUS1 | GDSL esterase/lipase | Cutin assembly | 1.95 | 6.57E-191 |
| Solyc01g088400 | CER1-like | CER1 Fatty acid hydroxylase | Wax synthesis | 1.05 | 5.52E-06 |
| Solyc03g005320 | KCS-like | Fatty acid elongase 3-ketoacyl-CoA synthase | Wax synthesis | 3.27 | 7.59E-05 |
| Solyc08g067260 | KCS10 | Fatty acid elongase 3-ketoacyl-CoA synthase | Wax synthesis | 1.08 | 9.75E-35 |
| Solyc11g067180 | CER4-like | Fatty acyl coA reductase | Wax synthesis | 5.99 | 2.44E-04 |
| **Down-regulated** |  |  |  |  |  |
| Solyc07g056320 | GPAT1 | ER glycerol-phosphate acyltransferase | Cutin synthesis | -2.40 | 1.17E-24 |
| Solyc03g065250 | CER1 | CER1 Fatty acid hydroxylase | Wax synthesis | -1.72 | 1.12E-22 |
| Solyc03g117800 | CER1-like | CER1 Fatty acid hydroxylase | Wax synthesis | -1.64 | 8.16E-28 |
| Solyc10g080840 | MAH1 | Cytochrome P450 | Wax synthesis | -2.48 | 8.41E-27 |
| Solyc09g083050 | KCS2 | Fatty acid elongase 3-ketoacyl-CoA synthase | Wax synthesis | -4.33 | 1.14E-65 |
| Solyc10g009240 | KCS1 | Fatty acid elongase 3-ketoacyl-CoA synthase | Wax synthesis | -1.35 | 2.43E-04 |
| Solyc06g074390 | CER4/FAR3 | Fatty acyl coA reductase | Wax synthesis | -3.19 | 3.74E-14 |
| Solyc09g092270 | CER2-like | Hydroxycinnamoyl-CoA shikimate/quinate hydroxycinnamoyl transferase | Wax synthesis | -1.59 | 8.39E-13 |

In order to assess the relationship between ABA signaling and cuticle formation, we examined the expression of ABA signaling core component genes in fruit skin of both *SlPP2C3*-RNAi lines and WT (Fig. 9C; Fig. 10C). Expressions of ABA receptor genes *SlPYL3* and *SlPYL8* were up regulated in transgenic lines. *SlPP2C1,* the only group A PP2C member, showed mild downregulated except for *SlPP2C3.* Among eight *SnRK2* members, expressions of *SlSnRK2.4, SlSnRK2.6* and *SlSnRK2.8* were significantly up regulated in transgenic lines. In addition, *SlPYL3* and *SlPP2C3* expressions were enriched in epidermis, and were co-expressed with *SlMYB96* and *SlMYB106* (Fig. S7). These data imply that the altered ABA signaling transduction in *SlPP2C3*-RNAi lines may be responsible for the changes of cuticle metabolism genes.

**Fig. 10.**
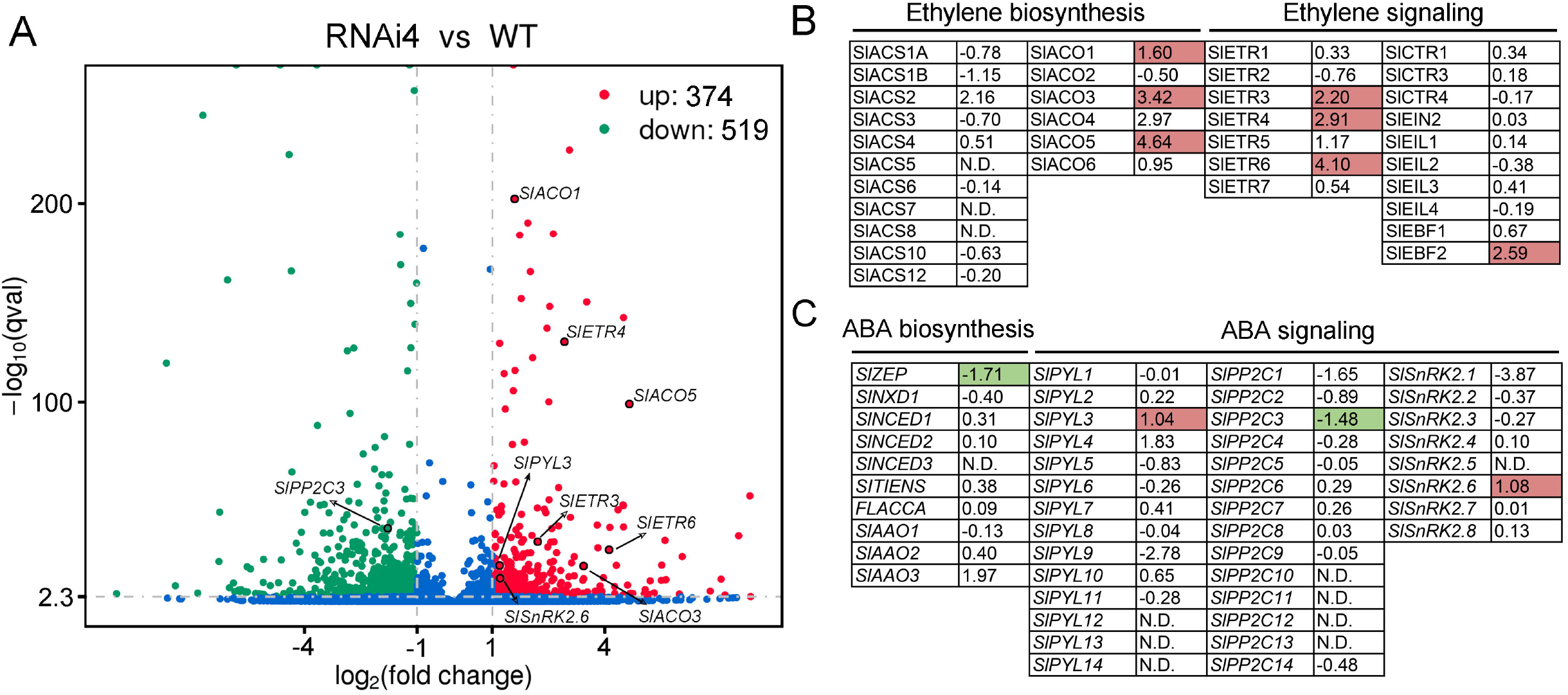
SlPP2C3 affects the transcription of genes involved in ethylene and ABA pathway. (A) Differential expressed genes between *SlPP2C3*-RNAi4 and WT fruits at mature green stage shown by volcano plot. (B, C) Log2 fold change values of genes involved in ethylene and ABA pathway in *SlPP2C3*-RNAi4 fruits compared to the WT. The statistically significant changes in color: red indicates up-regulation, green indicates down-regulation. Fold change > |1|, adjusted *p* value < 0.001, N.D., not detectable.

## Discussion

### Preferential interaction of SlPP2C3 with monomeric receptors

ABA initiates signal transduction by forming ABA-PYL-PP2C ternary complex. Both PYLs and PP2Cs are multigene families with redundant function. According to oligomeric state, PYL receptors were divided into dimeric receptor and monomeric receptor (Hao *et al.*, 2011). Group A PP2Cs were clustered into two subfamilies (Fig. S3) which determined the complexity of PYL-PP2C interaction. In our data, SlPP2C3 showed no interaction with dimeric receptors SlPYL5 and SlPYL8 in both yeast and plant cell. Similar result was obtained in our previous work as well (Chen *et al.*, 2016). Phylogenetic analyses indicated that SlPP2C3 is the ortholog of *Arabidopsis* AtHAI1/2/3. In *Arabidopsis,* HAI PP2Cs have marked preference for interactions with monomeric PYLs and showed no interaction with dimeric receptors AtPYR1, AtPYL1 and AtPYL2 (Bhaskara *et al.*, 2012). However, Antoni et al. showed that AtHAI1 activity could be inhibited by dimeric receptors *in vitro* (Antoni *et al.*, 2012). The molecular mechanism of the selective interaction between HAI-like PP2Cs and dimeric receptors is unknown. In our data, by generating a series mutant on PP2C-PYL interaction key residues SlPP2C3, we identified Arg-285 and Ser-295 might play an important role in preventing SlPP2C3 from interaction with dimeric receptors (Fig. 3). This result partially explain the mechanism of PP2C preference. However, more evidence from other members of subfamily II PP2Cs are required.

### SlPP2C3 negatively regulates ABA signaling and affects fruit ripening

In our study, SlPP2C3 acts as a negative regulator in tomato ABA signaling which was supported by two pieces of evidence: (1) SlPP2C3 interact with multiple SlPYLs and SlSnRK2.8 (Fig. 1; Fig. 2A) and (2) *SlPP2C3*-RNAi plants were hypersensitive to ABA while in *SlPP2C3-OE* plants, ABA were less sensitive in several ABA mediated processes (Fig. 5; Fig. 6). SlPP2C3 acts negatively in ABA signaling which is similar to other group A PP2Cs reported in several plant species. (Gosti *et al.*, 1999; Merlot *et al.*, 2001; Saez *et al.*, 2006; Rubio *et al.*, 2009; Li *et al.*, 2015; Zhang *et al.*, 2018).

It is worthy to notice that the fruit ripening schedule was affected by *SlPP2C3* expression (Fig. 7A, B). Previous studies demonstrated the role of endogenous ABA level and exogenous ABA treatment on tomato fruit ripening (Zhang *et al.*, 2009; Sun *et al.*, 2012b; Sun *et al.*, 2017; Liang *et al.*, 2020). More recent studies indicate that ABA signaling also plays a crucial role in regulating fruit ripening process. For example, the overexpression of tomato ABA receptor *SlPYL9* or suppression of *SlPP2C1* causes early fruit ripening (Zhang *et al.*, 2018; Kai *et al.*, 2019). In addition, ABA signaling inhibitor AA1 inhibits tomato fruit ripening (Ye *et al.*, 2017). Our data indicates that tomato fruit ethylene production is affected by SlPP2C3 (Fig. 7C) and several studies show the cross talk between PP2C and ethylene production. For instance, in *Arabidopsis* AtABI1 interacts with AtACS2 or AtACS6 and inhibits their activity by dephosphorylation (Ludwikow *et al.*, 2014). ABA inhibits ethylene production through ABI4-mediated transcriptional repression of AtACS4 and AtACS8 (Dong *et al.*, 2015). In our transcription data, several genes are involved in ethylene biosynthesis *(SlACOs)* and signaling *(SlETRs)* was up-regulated in *SlPP2C3*-RNAi fruit (Fig. S6). Taken together, we propose that SlPP2C3 negatively affects fruit ripening via the suppression of ethylene production.

### SlPP2C3 affects fruit outer epidermis cuticle formation

Fruit outer epidermis is the first barrier protecting fruit from desiccation and pathogen invasion, and it also plays a crucial role in fruit shelf life. ABA plays a role in cuticle formation and cuticle associated gene expressions under water deficit condition in *Arabidopsis* and tomato (Kosma *et al.*, 2009; Curvers *et al.*, 2010; Wang *et al.*, 2011). Recent study indicated that ABA core signal affected cuticle development in *Arabidopsis* not through transcription factors ABF2/3/4, which are the direct targets of SnRK2. This result implies the existence of a divergent pathway in the down-stream of SnRK2 kinase (Cui *et al.*, 2016). Martin et al. reported that tomato ABA deficient mutants exhibited reduced cutin and wax composition and down-regulated cuticle biosynthesis genes in leaf. However, ABA deficiency distinctly effected tomato fruit cuticle compared to leaf (Martin *et al.*, 2017). In our study, *SlPP2C3*-RNAi fruits exhibit altered epidermis surface (Fig. 9A), and further transcriptome data identified 2 MYB *(SlMYB96* and *SlMYB106)* transcription factor and 25 cuticle metabolism genes which were differentially expressed in *SlPP2C3*-RNAi fruit skin (Table 1). These MYB transcription factor may be the potential down-steam targets in ABA signaling pathway. Considering the similar expression patterns of *MYB96, MYB106, SlPYL3* and *SlPP2C3* as well as the unique localization of SlPYL3-SlPP2C3 complex (Fig. 2A; Fig. S8), we speculated that SlPYL3-SlPP2C3 mediated up-stream ABA signaling may be important in cuticle formation regulation. However, the precise mechanism remains open.

## Supplementary data

Fig. S1. Phylogenetic analysis of tomato and *Arabidopsis* type 2C protein phosphatase (PP2C) family members.

Fig. S2. Alignment of the amino acid sequences of tomato and *Arabidopsis* group A PP2C members.

Fig. S3. Phylogenetic analysis of tomato and *Arabidopsis* group A PP2C family members.

Fig. S4. Expressions of *SlPP2C3* in *SlPP2C3* transgenic lines.

Fig. S5. Vegetative phenotypes of *SlPP2C3* transgenic lines.

Fig. S6. Expression levels of genes related to ethylene synthesis and signaling in *SlPP2C3*-RNAi and WT fruits.

Fig. S7. Carotenoids, sugar and titratable acid in WT and *SlPP2C3* transgenic fruits.

Fig. S8. Spatiotemporal expressions of *SlPP2C3, SlPYL3, SlMYB96* and *SlMYB106* during M82 tomato fruit development and ripening.

Table S1. Basic information of tomato group A *PP2Cs* genes.

Table S2. Primers used for *SlPP2C*3-OE and *SlPP2C3*-RNAi plasmid construction.

Table S3. Primers used for bimolecular fluorescence complementation and subcellular localization construct.

Table S4. Primers used for yeast two-hybrid assay.

Table S5. Primers used for site-directed mutagenesis.

Table S6. Primers used for qRT-PCR analysis.

Dataset 1. Total genes in the WT and *SlPP2C3*-RNAi fruit skin at mature green stage detected by RNA-Seq.

Dataset 2. Genes expressed differentially in *SlPP2C3*-RNAi fruit skin compared to WT by RNA-Seq.

## Acknowledgments

This work was financially supported by Israel Science Foundation (ISF)–National Natural Science Foundation of China (NSFC) Joint Scientific Research Program [grant no. 31661143046] and the NSFC (grant nos. 31772270 and 31572095).

